# Training enhances the ability of listeners to exploit visual information for auditory scene analysis

**DOI:** 10.1101/295766

**Authors:** Huriye Atilgan, Jennifer K. Bizley

## Abstract

The ability to use temporal relationships between cross-modal cues facilitates perception and behavior. Previously we observed that temporally correlated changes in the size of a visual stimulus and the intensity in an auditory stimulus influenced the ability of listeners to perform an auditory selective attention task (Maddox et al., 2015). In this task participants detected timbral changes in a target sound while ignoring those in a simultaneously presented masker. When the visual stimulus was temporally coherent with the target sound, performance was significantly better than when it was temporally coherent with the masker sound, despite the visual stimulus conveying no task-relevant information. Here, we trained observers to detect audiovisual temporal coherence and asked whether this improved their ability to benefit from visual cues during the auditory selective attention task. We observed these listeners improved performance in the auditory selective attention task and changed the way in which they benefited from a visual stimulus: after training performance was better when the visual stimulus was temporally coherent with either the target or the masker stream, relative to the condition in which the visual stimulus was coherent with neither auditory stream. A second group which trained to discriminate modulation rate differences between temporally coherent audiovisual streams improved task performance, but did not change the way in which they used visual information. A control group did not change their performance between pretest and post-test. These results provide insights into how crossmodal experience may optimize multisensory integration.

## Introduction

Building an accurate perception of the world around us requires that the brain appropriately link signals across modalities (Kayser & Shams, 2015). Auditory and visual signals arrive and are processed with different latencies. Consequently, cross modal signals can be perceived as synchronous across a range of onset asynchronies – known as the temporal binding window (Dixon & Spitz, 1980; Meredith, Nemitz, & Stein, 1987). Previous studies have demonstrated that there is short term plasticity in this window (Megevand, Molholm, Nayak, & Foxe, 2013; Navarra et al., 2005; Schormans & Allman, 2018; Vroomen, Keetels, de Gelder, & Bertelson, 2004; Zmigrod & Zmigrod, 2015) and that experience and longer-term training can narrow this window such that listeners more accurately judge synchronous from asynchronous stimuli (Bidelman, 2016; Dixon & Spitz, 1980; H. Lee & Noppeney, 2011; Powers, Hillock, & Wallace, 2009).

Training listeners to refine their temporal binding window has been observed to have varied consequences for multisensory integration. For example, training which was effective in narrowing the temporal window for spatial ventriloquism also decreased the likelihood of cross-modal interactions as indexed by spatial ventriloquism (McGovern, Roudaia, Newell, & Roach, 2016). In contrast, training improved visual temporal discrimination abilities, but did not influence the likelihood of perceiving sound induced flash illusions (Ryan A. Stevenson, Wilson, Powers, & Wallace, 2013). Finally, in another study in which listeners were trained to discriminate asynchronous from synchronous stimuli listeners were subsequently shown to have stronger spatial ventriloquism effects when auditory-visual signals were temporally synchronous but spatially separated (Sürig, Bottari, & Röder, 2018).

A problem in interpreting these varied effects is that many lab-based tasks do not well capture the complexity that the brain faces in real-world situations. In most lab-based paradigms observers often judge single audio and visual signals, presented in an otherwise quiet and dark environment. In contrast, in the world, brain must match one of several competing sounds to a given visual object (or vice versa). Moreover, due to the variance in the timing of real-world signals, simply narrowing the window over which integration occurs may be a suboptimal strategy for effective information integration. Rather, what observers need to do is detect whether temporal coherence exists between signals in different modalities so that they may be appropriately grouped (Lee, Maddox, & Bizley, 2019).

An additional consideration for training studies that focus on the temporal binding window is that it is unclear to what extent any adjustments in the temporal binding window extend to other multisensory processing tasks. In many cases the task used to train observers is the same one used to measure the temporal binding window raising the question of how generalizable results are and whether they represent a genuine change in cross-modal binding, or whether listeners are simply shifting an internal criterion in order to improve their performance in this one task (Bizley, Maddox, & Lee, 2016).

In this study our goal was to ask whether we could train listeners to improve their ability to detect audiovisual temporal coherence and, in doing so, whether this would improve their ability to use visual cues to appropriately group sound elements from one stream and separate them from those elements in a competing sound. To train listeners to detect audiovisual temporal coherence listeners were required to differentiate streams in which audio and visual stimuli were amplitude / radius modulated in a statistically independent manner from stimuli in which audio and visual elements maintained some degree of temporal coherence. We elected to train listeners to detect small amounts of correspondence (rather than detect incoherence) as we reasoned that detecting moments of genuine correspondence is more likely to be useful than detecting incoherence.

In addition to measuring the ability of observers to assess temporal coherence before and after training (using similar stimuli to those used in the training), we used the auditory selective attention task from Maddox et al., (2015) to assess how effectively listeners could utilize visual information during the performance of an auditory selective attention task. This task required that subjects focused on one of two competing auditory streams and reported brief timbre perturbations within the target stream. They also watched a visual stimulus whose radius could change in a manner that was temporally coherent with either the target, the masker or neither auditory stream (but was never predictive about the timing of the timbre perturbations). In Maddox et al., we reported that the visual coherence condition significantly influenced performance in the auditory selective attention task such that performance was better when the visual stimulus was coherent with the target masker stream than when it was coherent with the masker stream, In order to determine whether any training effects we observed were critically dependent on improved temporal coherence detection, as opposed to passive exposure to temporally coherent auditory-visual streams, we also trained another group of observers in an amplitude modulation rate discrimination task with the same temporally coherent stimuli. A control group simply performed the pretest and post-test without any training. We hypothesized that an improved ability to detect temporal coherence may enable listeners to appropriately group temporally coherent audiovisual streams which in turn would promote more effective auditory selective attention. Our results support this hypothesis and demonstrate that only listeners trained to detect audiovisual temporal coherence change the way in which they are able to use visual information to augment auditory scene analysis.

## Methods

### Subjects

42 adults (age range 18–34 years; mean age 28 years; 11 males) with normal hearing and normal or corrected-to-normal vision, participated in the study. Six participants were excluded after the pretest due to poor performance (mean d’<0.8, n = 4), or low visual hit rates (<70%, n=2). The remaining 36 participants were included for further analysis and were randomly allocated to 3 groups (12 listeners per group). The study was approved by the Ethics Committee of the University College London (ref: 5139) and all procedures performed were in accordance with the Declaration of Helsinki. All individuals were paid for their participation and signed an informed consent form before participation.

### Stimuli and testing procedure

Each participant performed a “pretest” and “post-test” that consisted of the auditory selective attention task and an AV coherence detection threshold test. The pretest and post-test sessions each lasted 90 minutes and were separated by a maximum of 2 weeks and a minimum of 5 days. Participants in the control group did no training sessions but performed the pretest and post-test within 2 weeks. Participants in the two training groups performed 5 training sessions on 5 separate days over not more than 2 weeks (Figure 1A).

**Figure 1:**
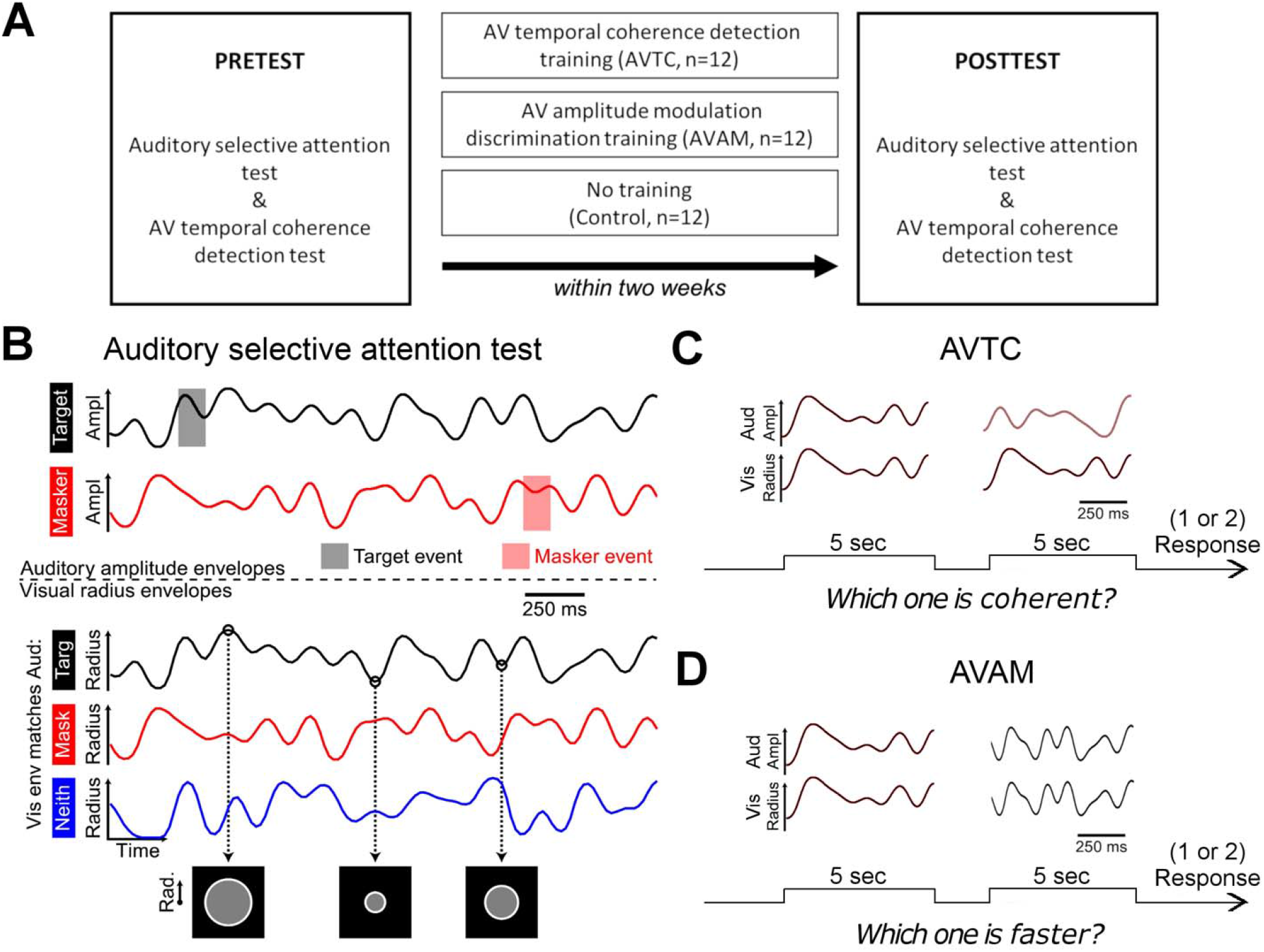
**A** Experimental design. **B** Schematic representation of auditory and visual stimuli used in the auditory selective attention test (panel was taken from Maddox et al., 2015). Amplitude envelopes of target (black) auditory stream and masker (red) auditory stream and visual radius envelopes for three auditory visual (AV) coherence conditions; target coherent (black), masker coherent (red) and neither (blue). Examples frames of the visual stimuli at three radius level. **C** Schematic representation of AV temporal coherence detection test/ AVTC training. Two 5 seconds AV pairs were used. One maintained some degree of temporal coherence (left, here fully coherent) while the other was always fully independent (right) **D** Schematic representation of AV amplitude modulation rate discrimination (AVAM) training. Two 5 second temporally coherent AV pairs were used. Each pair had a different modulation envelope (and rate) but stimuli were always temporally coherent across modalities.

#### Auditory selective attention task

The auditory selective attention task required that listeners attend to one of two competing auditory streams and report the presence of brief (200 ms) timbre perturbations in the target audio stream. They were additionally required to monitor a visual stimulus whose radius changed in time. The two audio streams were independently amplitude modulated and the visual radius was modulated with a time course that matched one or the other auditory stream or was independent of them both (Figure 1B). Envelopes for visual envelope and auditory amplitude were created using the same frequency domain synthesis. For each trial, an envelope was created by first setting all amplitudes of frequency bins above 0 Hz and below 7 Hz to unity and others to zero. At an audio sampling rate of 24,414 Hz, all non-zero bins were given a random phase from a uniform distribution between 0 and 2π, the corresponding frequency bins across Nyquist frequency were set to the complex conjugates to maintain Hermitian symmetry, and the inverse Fourier transform was computed yielding a time domain envelope. Second and third envelopes were created using the same method, and orthogonalized using a Gram-Schmidt procedure. Each envelope was then normalized so that it spanned the interval [0, 1] and then sine-transformed [y = sin2 (πx/2)] so that the extremes were slightly accentuated. Visual envelopes were created by subsampling the auditory envelope at the monitor frame-rate of 60 Hz, starting with the first auditory sample, so that auditory amplitude corresponded with the disc radius at the beginning of each frame.

Stimuli were presented in an unlit sound-attenuating room over headphones (HD 555, Sennheiser, Wedemark, Germany). Subjects were seated 60 cm from the screen with their heads held stationary by a chinrest. Auditory stimuli were created in MATLAB and presented using an RP2 signal processor (Tucker–Davis Technologies, Alachua, FL, USA). Each began and ended with a 10 ms cosine ramp. All stimuli were presented diotically. Visual stimuli were synthesized in MATLAB (The Mathworks, Natick, MA, USA) and presented using the Psychophysics Toolbox (Brainard, 1997). The visual stimuli were gray discs that subtended between 1° and 2.5°. The white ring extended 0.125° beyond the gray disc.

The auditory stimuli were generated as described in the *timbre* variant of Maddox et al., (2015). Each auditory stream was generated as a periodic impulse train and then filtered with synthetic vowels simulated as four-pole filters (formants F1–F4). The /u/ stream had formant peaks F1–F4 at 460, 1105, 2857, 4205 Hz and moved slightly towards /ε/ during timbre events, with formant peaks at 730, 2058, 2857, 4205 Hz. The /a/ stream had formant peaks F1–F4 at 936, 1551, 2975, 4263 Hz and moved slightly towards /i/ during timbre events, with formant peaks at 437, 2761, 2975, 4263 Hz. During timbre events the formants moved linearly toward the deviant for 100 ms and then linearly back for 100 ms. Streams were calibrated to be 65 dB SPL (RMS normalized) using an artificial ear (Brüel & Kjær, Nærum, Denmark) and presented against a low level of background noise (54 dB SPL). Unlike Maddox et al., we did not assess individual timbre discrimination thresholds but instead used a fixed level of difficulty determined using the average of the individual thresholds measured previously. For [e] deviants in [u] stimuli this corresponded to a shift of 42 Hz in F1 frequency and 143 Hz for F2, and for [i] deviants in [a] stream there was a shift of 75 Hz for F1, 196 Hz for F2. Both vowels could take either F0 value (175 Hz or 195 Hz, counterbalanced) and were equally likely to be target or masker. Trials lasted 14 seconds. They began with only the target auditory stimulus and the visual stimulus, indicating the to-be-attended (target) auditory stream to the subject. The to-be-ignored auditory stream (masker) began 1 second later. As with the rest of the trial, the visual stimulus was only coherent with the auditory target during the first second if it was a match-target trial. All streams ended simultaneously. Events did not occur in the first two seconds (i.e. 1 second after the masker began) or the last 1 second of each trial, or within 1.2 s of any other events in either modality. A response made within 1 second following an event was attributed to that event. To ensure audibility and equivalent target to masker ratios without providing confounding information to the subject, an event in either auditory stream or the visual stream could only begin when both auditory envelopes were above 70% maximum. There were between 1 and 3 inclusive events (mean events = 2) in both the target and masker in each trial. There were also between 0 and 2 inclusive visual flashes per trial (mean flashes = 1), in which the outer ring changed from white to cyan (0% red, 100% blue, 100% green) and back. Each subject completed 32 trials of each temporal coherence conditions (96 totals), leading to 64 potential hits and 64 potential false alarms for each condition (i.e., 128 responses considered for each d′ calculation) as well as 32 visual flashes per condition. When computing d′, auditory hit and false alarm rates were calculated by adding 0.5 to the numerator and 1 to the denominator so that d′ had finite limits.

#### Auditory-visual temporal coherence detection test

The AV temporal coherence detection test was used to determine perceptual thresholds for detecting AV temporal coherence. Auditory and visual stimuli were generated as described in the auditory selective attention task above, with a single auditory and visual stream presented on each occasion. The pitch and timbre of stimuli were varied across trials such that the auditory stimuli were either [u] or [e] (without any timbre deviants embedded), with F0 = 175 or 195 Hz, counterbalanced. The sounds in both stimulus intervals within the trial had identical pitch and timbre values and had a duration of 5 seconds. In one stimulus interval, the sound was accompanied by a visual stimulus in which the radius changed over time independently of the auditory stimulus. In the other intervals, the auditory and visual stimulus maintained some degree of temporal coherence (Figure 1C). The method of constant limits was used to determine the threshold with subjects performing 20 trials at each coherence level. AV stimuli were generated from 100% coherent in 10% steps to 10% coherent by multiplying the temporally coherent envelope with an independent envelope. Participants were required to select the interval (by pressing 1 or 2 on a button box) in which the temporally coherent pair was presented. Feedback was provided on every trial.

#### Auditory-visual temporal coherence (AVTC) training

The stimuli and procedure in the AVTC training were identical to those used in the threshold test, but with an adaptive three-down one-up rule to determine the coherence level of the stimulus in the next trial. Previous work has demonstrated that task difficulty is an important aspect in driving multisensory learning (De Niear, Koo, & Wallace, 2016) so by using this approach we required that participants worked near to their threshold for a large proportion of the training session. In the first training session, the stimuli in the first trial were 100% coherent, and 100% independent. For the first 6 reversals coherency was decreased in 10% steps followed by 5% steps for the following six reversals and by 2.5% steps for the remainder. The procedure was terminated at 18 reversals unless a maximum of 150 trials was reached first. For the 2nd-5th training session the first “coherent” stimulus was generated with the average coherence level of the last ten reversals in the previous session. Each training session lasted less than 40 minutes. Feedback was provided on every trial. Participants performed 5 training sessions on separate days over not more than 14 days.

#### Amplitude modulation rate discrimination (AVAM) training

In AVAM training, two temporally coherent AV stimuli (5 seconds long, with a constant pitch and timbre within a trial, counterbalanced across trials as described in “auditory-visual temporal coherence detection test”, Figure 1D) were sequentially presented. In one interval, the envelope was always generated with a 7 Hz cut off rate, whereas the other was generated with a higher rate (maximum AM cut off rate = 11 Hz). In the sessions of AM rate training, an adaptive three-down one-up rule was used to determine the AM rate of the stimulus in the next trial. In the first session, the first stimulus was generated at the maximum AM rate and differed in AM rate by 1Hz for the first six reversals and 0.5 Hz for the next six reversals and 0.25Hz for the rest of the trials. The procedure was terminated at 18 reversals unless a maximum of 150 trials was reached first. In each consequent session the first stimulus was generated with the average coherence level of the last ten reversals in the previous session. Participants pressed “1” or “2” on the press box to indicate the interval of the faster AV pair. Feedback was provided on each trial.

### Statistical Analysis

Statistics were performed using MATLAB (2011b, Mathworks, USA) and SPSS (IBM). The d’, hit rates, false alarm, and visual hit rates across AV coherence conditions and pretest versus post-test were calculated and statistical significance across groups was assessed by two-tailed unpaired Student’s t-tests, Mann-Whitney U-tests, one-way analysis of variance (ANOVA), mix ANOVA or repeated measures of ANOVA where appropriate. Mauchly’s test of sphericity was used for the assumption of homogeneity of variance for independent tests in repeated measured analysis if not otherwise reported. Significant main effects or interactions were followed up with post-hoc testing using Bonferroni corrections where applicable. Significance was declared at p <◻0.05, with a precise p value stated in each case, and all tests were two-sided. Partial eta squared was reported for different ANOVA procedures in SPSS to demonstrate the effect size we observed. For individual differences in AV temporal coherence detection thresholds, 95% confidence intervals were calculated with a linear regression.

## Results

We recruited participants and randomly assigned them to one of three groups, each of which performed a pretest and a post-test. Pretests and post-tests comprised the timbre variant of the selective attention task in Maddox et al., (2015) and an AV temporal coherence detection threshold test. In between the pretest and post-test, one group trained on an AV temporal coherence task (AVTC group, n = 12), one group trained on an AM rate discrimination task using temporally coherent AV stimuli (AVAM group, n = 12), and a third group simply performed the pretest and post-test separated by a minimum of 5 days (control, n = 12; Figure 1A).

We first confirmed that both trained groups improved their ability on the trained stimulus feature. Figure 2A and B shows the thresholds derived from the last 5 reversals for the training session on day 1 and day 5 for the AVTC group and the AVAM group respectively. For both groups’ thresholds were significantly lower for session 5, than for session 1 (pairwise t-test on S1 and S5 AVTC thresholds, t_11_ = 2.961, p = 0.007; AVAM thresholds: t_11_ = 4.529, p<0.001).

**Figure 2:**
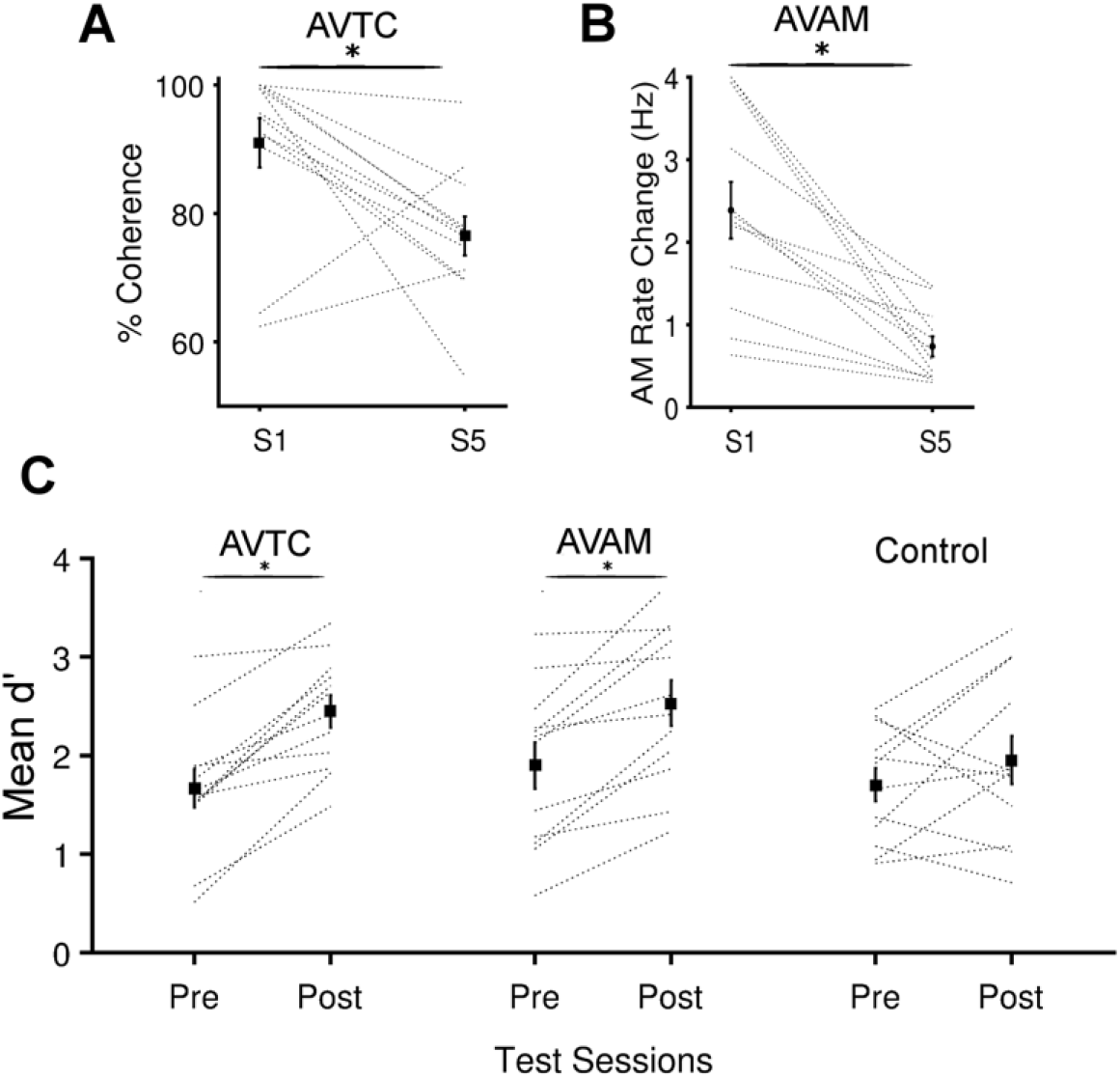
Training improved performance in trained tasks and the Auditory Selective Attention task. **A** Training in auditory visual temporal coherence detection (AVTC) task was effective at driving an improvement in coherence detection threshold between session (S1) and session 5 (S5). Black vertical lines show the mean ± SEM across participants, gray lines are individual subjects. **B** Training in the auditory visual amplitude modulation (AVAM) task was effective at driving an improvement in AM rate discrimination between S1 and S5. **C** Both trained groups improved their overall d’ in the auditory selective attention task. Mean d’ across AV coherence condition of pretest and post-test for the three groups. *indicates significant differences (Pairwise t-tests p < 0.05)

Next, we asked whether there was a change in performance in the auditory selective attention task between the pretest and the post-test. We anticipated that participants improve their performance between the pretest and the post-test in both trained groups, as these listeners had been exposed to the vowel sounds during their training sessions (although the sounds that they heard did not contain any timbre deviants).

To detect overall changes in performance that occurred independently of the AV coherence condition, we calculated the overall sensitivity (d’), and directly compared the pretest and post-test data across the three experimental groups (Figure 2C). We ran an 3×3×2 mixed ANOVA for *d*′, with a between-subjects factor of experimental groups (AVTC, AVAM and control) and within-subjects factors of sessions(pretest and post-test) and audio–visual coherence condition (target coherent, masker coherent, neither, data available as source data 1).

We found a significant effect of session (F (1, 33) = 35.411, p <0.001, partial η^2^ = 0.518), with no significant effect of experimental group (F (2, 33) = 1.030, p = 0.368, partial η^2^ = 0.059) and no significant interaction (F (2, 33) = 2.858, p = 0.072, partial η^2^ = 0.148). Post-hoc pairwise comparisons between pretest and post-test data revealed that means of d’ scores were significantly different for both trained groups (AVTC group: t_11_ = 3.065, p = 0.006, AVAM group: t_11_ = 1.920, p = 0.034; control group: t_11_ = 0.854, p = 0.402). Thus, both experimental groups, but not the control group, improved their ability to detect timbre deviants in a sound mixture.

We found a significant effect of coherence condition (F (2, 66) = 13.859, p <0.001, partial η^2^ = 0.296) and significant interaction between session and coherence condition (F (2, 66) = 3.302, p = 0.043, partial η^2^ = 0.091). In Maddox et al., (2015) we observed that performance in the auditory selective attention task varied with AV coherence condition, with performance when the visual condition was coherent with the target stream being superior to when the visual condition was coherent with the masker stream. Focusing on the pretest data obtained from naïve listeners, Bonferroni post-hoc tests showed that participants detect deviants significantly better in only target coherent condition than masker coherent condition, thus replicating our original observations.

Having observed that training increased performance on the auditory selective attention task, and that prior to training listeners showed a characteristic pattern of performance across AV coherence conditions, we went on to explore in more detail the effect of training on the way in which the visual condition influenced performance in the auditory selective attention task. To do this, we considered the d’ values across condition, and, for visualization purposes, calculated the normalized d’ values (note: all statistical comparisons are performed on the untransformed values). Normalised d’ illustrates how performance varies across visual condition by taking the difference between each condition and the across-condition mean. We hypothesized that if training had no influence on the way in which listeners were influenced by the coherence condition we would see the same across-coherence-condition pattern, simply shifted up to higher d’ values. In this situation the normalized d’ values would be unchanged by training indicating that the increase in d’ was uniform across coherence condition. In contrast, if there was a change in the magnitude of the visual stimulus induced effects we predicted a similar pattern of across-coherence-condition performance, but a larger difference between target and masker coherent conditions. This would be reflected in normalized d’ measures being larger in magnitude, but equivalent in sign between pretest and post-test. Finally, if training influenced the way in which listeners utilized visual information, we predicted that there would be a change in the pattern of across-coherence-condition d’ values, and a change in the sign of the normalized values.

### AVTC but not AVAM training alters the way listeners use visual cues in selective attention task

We therefore calculated d’, hit rates, false alarm rates and bias for each coherence condition in the auditory selective attention task, for each training group and the pretest and post-test separately. To analyse changes between pretest scores and post-test scores for each group, we performed a two-way repeated measure within-subjects ANOVA with factors of coherence condition (target coherent, masker coherent and neither) and session (pretest and post-test; Table 1). In this framework, we expect a significant effect of coherence condition in all groups and of training in the AVTC and AVAM group. Of particular interest is the interaction term, as this would indicate a training-induced change in the way in which visual information impacted performance.

**Table 1:** The results of two-way repeated measures within-subjects ANOVA for each variable (p < 0.05 in bold) for d’, hit rates, false alarm and bias for three experimental groups.

|  | Coherence conditions |  |  | Session<br>(pre vs post) |  |  | Interaction between coh.<br>condition & session |  |  |
| --- | --- | --- | --- | --- | --- | --- | --- | --- | --- |
| | F | p | $\eta^2$ | F | p | $\eta^2$ | F | p | $\eta^2$ |
| AVTC Group |  |  |  |  |  |  |  |  |  |
| $d'$ | 6.908 | **.050** | .386 | 36.245 | **<.001** | .767 | 7.258 | **.002** | **.493** |
| *Hit Rates* | 5.080 | **.015** | .316 | 7.731 | **.018** | .413 | 4.660 | **.021** | **.298** |
| *False Alarm\** | 2.692 | .115 | .197 | 17.164 | **.002** | .609 | 2.555 | .100 | .189 |
| *Bias* | 1.233 | .311 | .101 | 0.128 | .727 | .110 | 3.005 | .070 | .215 |
| AVAM Group |  |  |  |  |  |  |  |  |  |
| $d'$ | 5.723 | **.010** | .342 | 23.134 | **.001** | .678 | .009 | .854 | .014 |
| *Hit Rates* | 2.952 | .073 | .212 | 24.148 | **<.001** | .687 | .386 | .682 | .042 |
| *False Alarm\** | 2.317 | .145 | .174 | 5.846 | **.034** | .347 | .376 | .655 | .033 |
| *Bias\** | 1.909 | .190 | .148 | 3.337 | .095 | .233 | .922 | .411 | .077 |
| Control |  |  |  |  |  |  |  |  |  |
| $d'$ | 4.653 | **.021** | .297 | 1.431 | .257 | .115 | .039 | .962 | .004 |
| *Hit Rates* | 0.998 | .385 | .083 | 1.180 | .301 | .097 | .065 | .929 | .006 |
| *False Alarm* | 3.023 | .069 | .216 | 0.088 | .772 | .008 | .253 | .779 | .022 |
| *Bias* | 0.301 | .743 | .027 | 0.084 | .777 | .008 | .043 | .958 | .004 |

\*Mauchly's test of sphericity was significant ( $<0.05$ ), Greenhouse-Geisser corrected values are reported.

In the AVTC group, participants trained on a task in which they were actively judging the temporal coherence of auditory and visual streams. Prior to training, participants showed the expected effect of coherence condition (i.e. only target coherent > masker coherent). However, after training a different pattern was observed: both target and masker coherent conditions were superior to the neither condition in which the visual stimulus changed independently (Figure 3A). Notably, the normalized d’ measures for the masker coherent condition changed from negative to positive in this group (Figure 3D). Consistent with these observations a repeated ANOVA analysis of d’ scores revealed significant effect of session (F (1, 11) = 36.245, p <0.001), a borderline effect of coherence condition (F (2, 22) = 6.908, p=0.05) and a significant interaction (Figure 3A, D, G; F (2, 22) = 7.258, p = 0.002; see also Table 1 which reports effect sizes). Post-hoc comparisons across AV coherence condition in the pretest data revealed that participants performed better when the visual stimulus was coherent with the target auditory stream versus the masker auditory stream (target coherent > masker coherent). In contrast, post-hoc comparisons of the post-test d’ scores revealed that, after training, performance was better when the visual stimulus was coherent with either the target or the masker stream than in the condition which coherent with neither (target coherent > neither, masker coherent > neither, Bonferoni corrected post-hoc comparison p<0.05).

**Figure 3:**
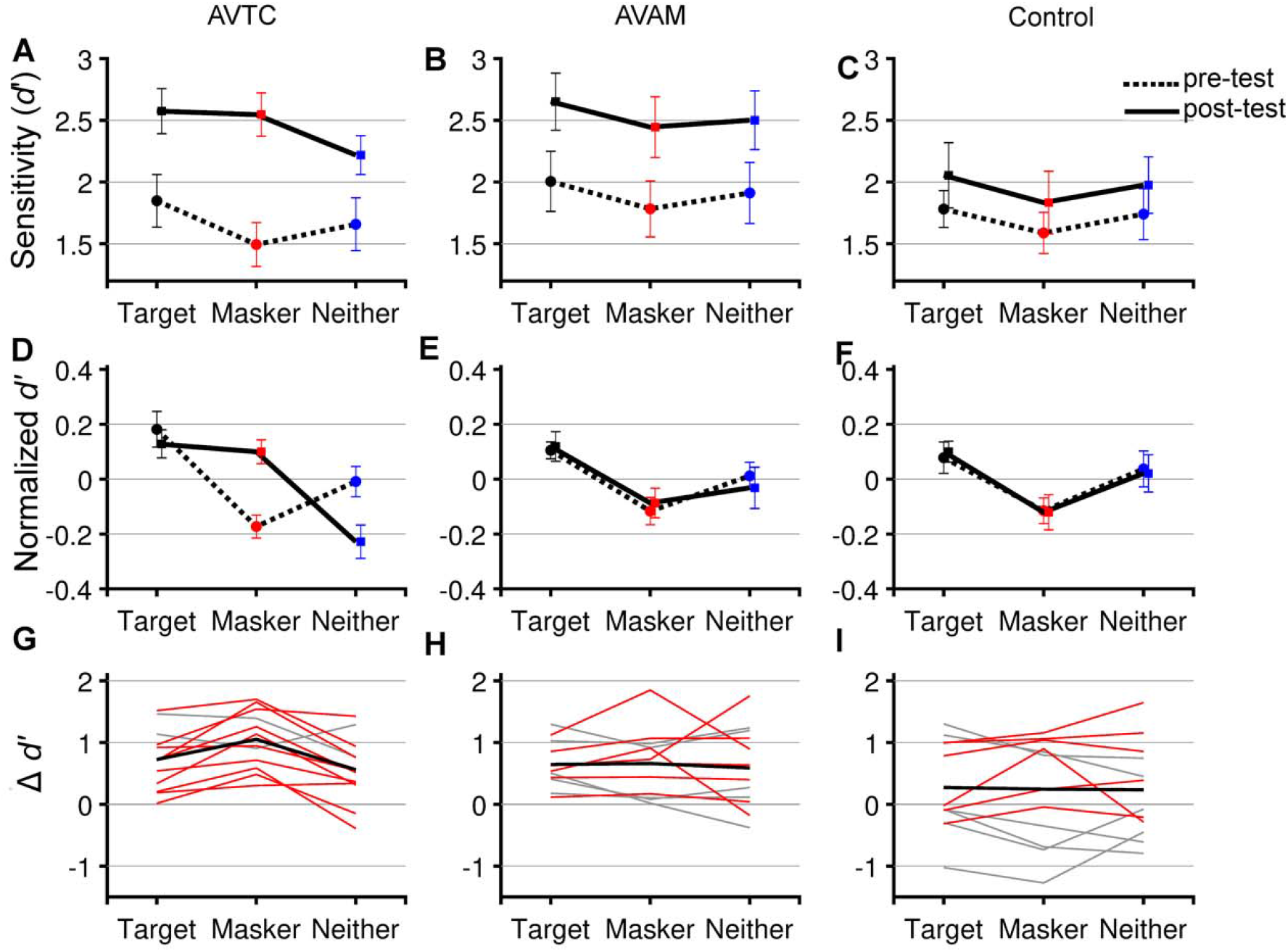
Training to detect AV temporal coherence changed how listeners utilized visual information. **A-C** pretest (dashed line) and post-test (solid line) performance in the auditory selective attention task according to AV coherence conditions. A, D, G: AVTC group, B, E, H: AVAM group, C, F, G: control group. **D-F** Normalized mean ± SEM performance (calculated as within condition d’ normalized to across condition performance for pretest and post-test separately). **G-I** the difference in d’ between pre and post-test for each participant in either gray, or red. Participants who showed a greater increase in the masker coherent condition than the target coherent after training are plotted in red, the group mean is plotted in bold black.

Participants in AVAM group were asked to detect the amplitude modulation rate of temporally coherent AV pairs. They were not actively detecting temporal coherence, but passively exposed to temporally coherent AV pairs. Although training improved their overall performance in the auditory selective attention task, the way in which they used visual information appeared unchanged after training. In both pretest and post-test performance was the best in the target coherent condition (Figure 3B) and the normalized d’ measures, which effectively factor out the overall d’ improvement, were overlapping (Figure 3E) suggesting that there was no change in the way in which the coherence condition effected performance (Figure 3H). Both session (F (1, 11) = 23.134, p = 0.001) and coherence condition (F (2, 22) = 5.723, p = 0.010) influenced d’, but – importantly – there was no interaction (F (2, 22) = 0.09, p = 0.854). Post-hoc comparisons (p<0.05) across coherence conditions revealed that subjects performed better when the visual stimulus was coherent with the target auditory stream compared to the masker coherent condition in both the pretest and post-test (target coherent > masker coherent). Therefore, this suggested an overall improvement in performance after AVAM training, but no change in the way in which subjects were able to exploit visual cues.

Performance in the control group did not differ significantly between pretest and post-test (Figure 3C, F, I): there was no effect of session (F (1,11) = 1.431, p = 0.257), but a significant effect of coherence condition (F (2, 22) = 4.653, p = 0.021), with no interaction (F (2, 22) = 0.039, p = 0.962). Participants performed significantly better when the visual stimulus was coherent with the target auditory stream versus the masker auditory stream (target coherent > masker coherent) in pretest and post-test (Figure 3F).

To better understand the effect in the AVTC group we considered the hit rates and false alarm rates (which together define d’) to determine whether the changes were principally driven by an improved ability to detect the target timbre deviations in the masker coherent condition, or an improved ability to ignore deviants that occurred in the masker stream. Figure 4 shows the changes in hit rates and false alarm rates between the pretest and post-test for the AVTC group and suggests that training drove an overall drop in false alarms across all coherence conditions, and a condition-specific increase in hit rates, with the largest increase occurring in the masker coherent condition. Two-way ANOVAs (table 1) revealed that there was a significant increase in hit rates with training, and a significant session (F (1, 11) = 7.731, p = 0.018), coherence condition (F (2, 22) = 5.080, p = 0.015), and interaction (F (2, 22) = 4.660, p = 0.021). The decrease in false alarm rate between pretest and post-test was also statistically significant (F (1,11) = 17.164, p = 0.002) without a significant coherence condition effect (F (2, 22) = 2.692, p = 0.115).

**Figure 4:**
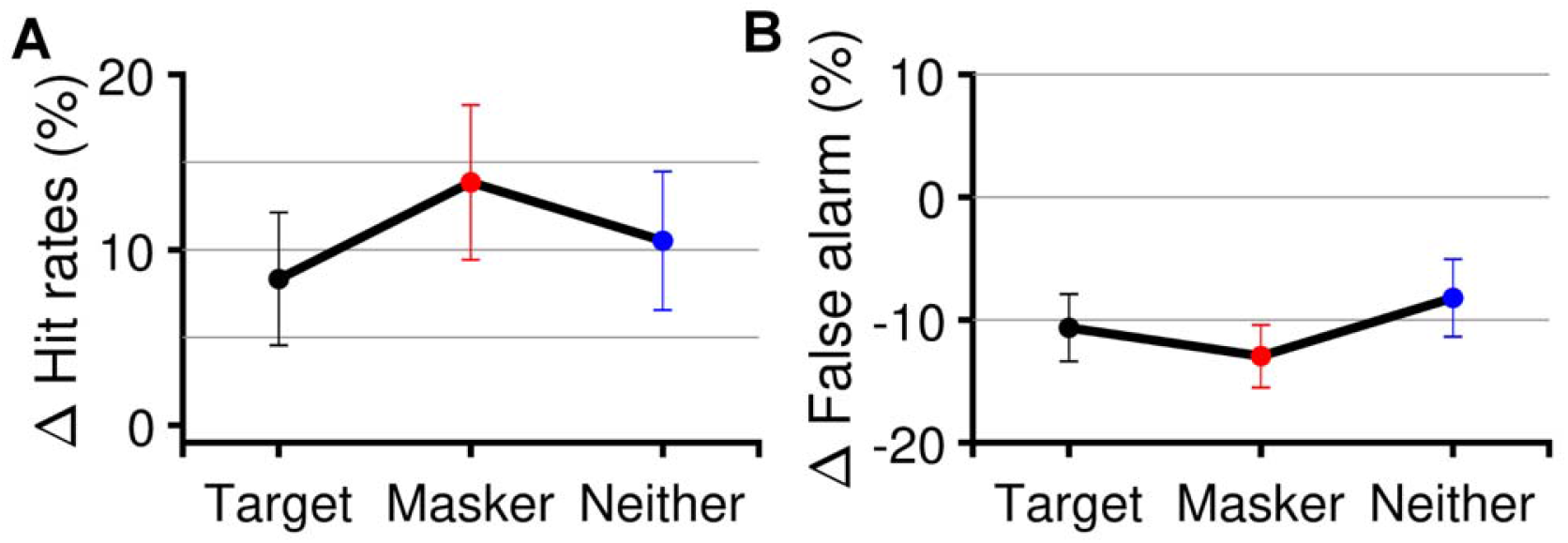
Improved performance in the masker coherent condition in AVTC group was driven by a drop in false alarms and an increase in the masker-coherent condition hit rate. The changes in hit rates (**A**) and false alarm rate (**B**) between pretest and post-test; mean ± SEM.

### AVTC training improved temporal coherence detection

The pretest and post-test included a temporal coherence detection threshold test for all listeners. As anticipated, those listeners trained to detect audiovisual temporal coherence improved their thresholds between the pretest and post-test: (Figure 5A, t_11_ = 3.081, p = 0.005). Listeners in the AVAM group who were exposed to temporally coherent AV stimuli did not improve their thresholds (t_11_ = 1.69, p = 0.104) or not in the control group (t_11_ = 0.234, p = 0.817). Examination of these data also revealed there was considerable variability in how well observers could detect temporal coherence, and in the AVAM group, considerable individual variability in the way in which performance changed between tests. We therefore asked whether any of this individual variability predicted performance in the auditory selective attention task.

**Figure 5:**
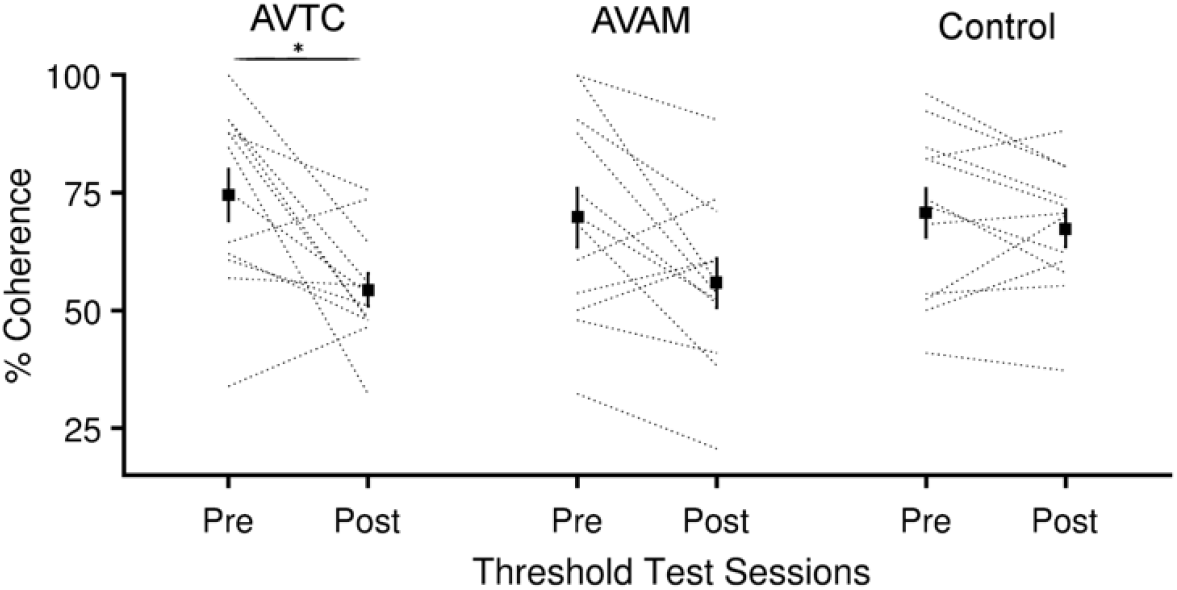
Training decreased temporal coherence thresholds in the AVTC group. AV temporal coherence threshold values of pretest and post-test for three groups. * indicates significant paired t-test comparison (p<0.05).

### Sensitivity to temporal coherence is not predictive of performance in naïve listeners

We first explored whether individual differences in temporal coherence detection ability accounted for the impact that the coherence condition had on performance in naïve listeners. Specifically, we tested the hypothesis that the ability of naïve listeners to detect AV temporal coherence would predict their ability to benefit from AV temporal coherence in the auditory selective attention task. We correlated each listener’s AV temporal coherence threshold with the difference between the d’ score in the target and masker coherent condition (Figure 6A). Contrary to this hypothesis, there was no relationship between these values (r = 0.1543, p = 0.3688), nor was there any relationship between overall performance (across condition d’) and AV temporal coherence thresholds (r = 0.2888, p = 0.0882).

**Figure 6.**
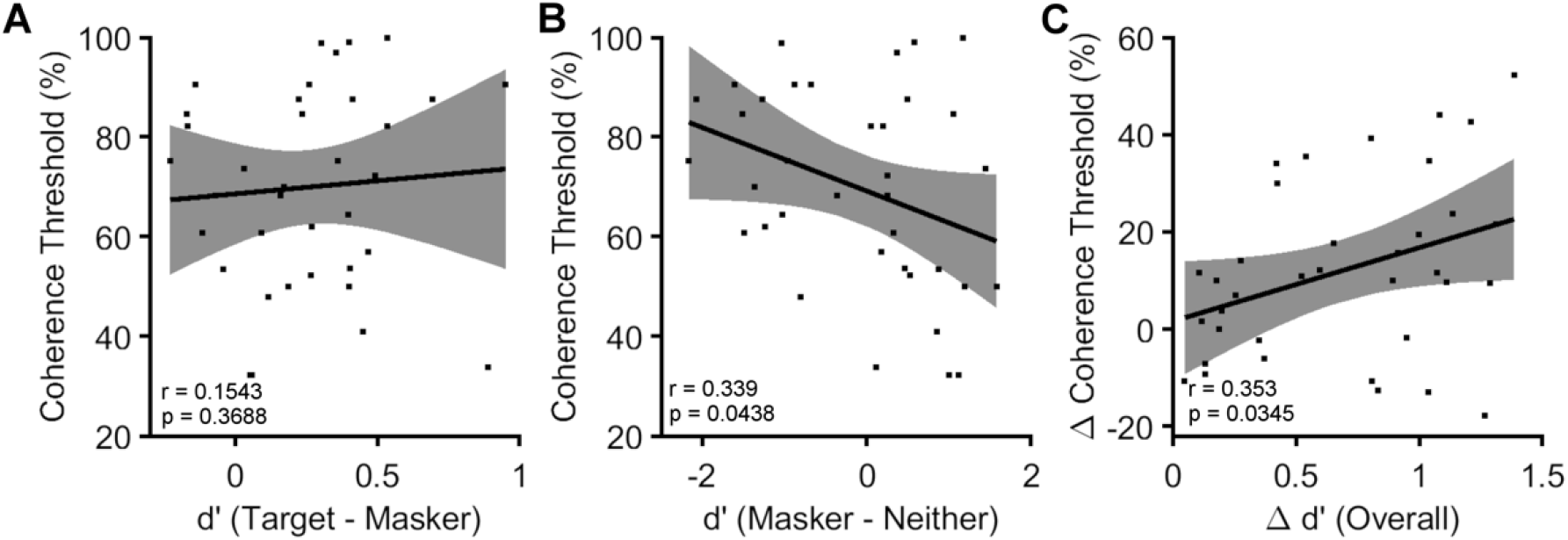
Individual differences in AV temporal coherence detection. **A** Target coherent -masker coherent d’ difference (n=36 naïve listeners) versus AV temporal coherence threshold (low values indicate better thresholds) from all listeners pretest data. Error bars indicate 95% confidence intervals **B** Masker-neither d’ difference versus AV coherence threshold for the pretest data. **C** The change in overall performance between the pretest and post-test versus change in AV coherence threshold.

Having observed that the AVTC group improved their ability to utilize audiovisual temporal coherence in the masker coherent condition, we considered whether temporal coherence thresholds might be correlated with the magnitude of the benefit / impairment that the masker coherent condition had over the neither condition. To assess this, we considered the difference in d’ between the masker coherent condition and the neither condition. This comparison was weakly negatively correlated with AV temporal coherence thresholds for naïve listeners (Figure 6B; r = 0.339, p = 0.0438) suggesting a trend where participants with better AV temporal coherence thresholds were more able to exploit the temporal coherence between masker stream and visual stimulus to yield a performance benefit relative to the neither condition. This finding mirrors the effect of training whereby improving AV coherence thresholds led to an improvement in the masker coherent condition.

Finally, since some participants in all groups showed improved temporal coherence detection thresholds, we asked whether at an individual subject level whether the change in temporal coherence detection threshold correlated with the overall change in d’: Participants with a larger change in their AV coherence threshold showed larger improvements in overall performance (Figure 4C; r = 0.353, p = 0.0347).

## Discussion

Here we demonstrate that five short training sessions can improve a listener’s ability to detect AV temporal correspondence and change the way in which they are able to exploit cross-modal temporal coherence. In naïve listeners a visual stimulus that is temporally coherent with a target auditory stream enhances performance relative to when the visual stimulus is temporally coherent with the masker stream (with a condition which was coherent with neither yielding intermediary performance). After training, both target and masker visual coherence conditions yielded significantly better performance relative to the condition in which the visual stimulus was coherent with neither.

We had two control groups in this study. The first did not perform any training in between the pretest and the post-test. The second group was trained on a temporal rate discrimination task that utilized temporally coherent auditory visual stimuli. Therefore, like the AVTC group they judged temporal features of the stimuli and were exposed to the stimulus streams that formed the target sounds in the auditory selective attention task. Unlike the AVTC group, the AVAM group did not require that observers make a cross-modal discrimination and observers were free to base their decisions on auditory and/or visual features. Consistent with perceptual learning resulting from exposure to the sounds, both groups improved in their performance of the auditory selective attention task. However, only observers that were required to explicitly judge cross-modal temporal coherence showed a change in the way in which visual information was used for auditory scene analysis.

We have previously argued (Maddox et al., 2015) that the visual stimulus impact performance in the auditory selective attention task by altering how well listeners are able to separate the two competing streams and select the target. This is because the features that link the audio and visual streams (temporally coherent changes in auditory intensity and visual size) are independent of the changes in sound timbre that listeners are required to detect and the visual stimulus itself conveys no information about whether or when (or in which stream) the auditory timbre deviants occurred. Thus, improved performance in the auditory task demonstrated that auditory and visual streams have been bound into a single perceptual object (Bizley et al., 2016; Lee et al., 2019). Recordings in the auditory cortex of passively listening, naïve ferrets demonstrate that audiovisual temporal coherence causes an enhanced representation of the temporally coherent stream that extends to all of its features (Atilgan et al., 2018). As well as providing further evidence for audiovisual object formation, assuming that an audiovisual object has processing advantages, or captures selective attention more strongly, these data provide a bottom-up explanation for how the enhanced performance in the target coherent condition and the impaired performance in the masker coherent condition arises. What therefore might be the mechanism through which visual coherence with a to-be-ignored sound yields a processing advantage?

At a cellular level, successful stream segregation is thought to be a consequence of the activation of distinct neural populations in time, such that neurons representing the same stream are highly temporally coherent in their responses and those representing different streams share low coherence (Lu et al., 2017; Middlebrooks & Bremen, 2013). Under such a model temporal coherence with either the target *or* the masker stream should result in more distinct (and hence better segregated) neural responses that in turn offer a more effective substrate on which selective attention can operate. If AV temporal coherence allows the representation of each of two competing sounds to be more distinct within sensory cortex then temporal coherence between target or masker stream should offer an advantage over an independently modulated visual stimulus. The data from the AVTC group suggest that after training listeners were able to benefit from AV temporal coherence when the visual stimulus was temporally coherent with either auditory stream. Possible mechanistic explanations for this would be that training has resulted in listeners being better able to use top-down control to actively suppress the masker stream in the masker coherent case, which in turn enables them to better detect the timbre deviants in the target stream. This suggestion is supported by the data in figure 4, which show a condition-specific increase in hit-rates for the masker-coherent condition after training. While we can only speculate about the mechanism underlying the effects observed here, a recent study that trained listeners to improve their audiovisual temporal perception reported enhanced beta band activity after training and suggested that enhanced top-down modulation was responsible for improved temporal processing (Theves, Chan, Naumer, & Kaiser, 2020). Further assessment of this hypothesis requires neurophysiological work to determine how selective attention and audiovisual object formation interact to shape the responses to target and masker streams in auditory cortex.

We implemented an adaptive training procedure with feedback to force subjects to work close to their perceptual threshold for most of the training periods. In keeping with other studies (Sürig et al., 2018), adaptive training was highly effective at rapidly driving learning in both of the temporal discrimination tasks. While many adaptive tasks show that the majority of learning occurs in the first session, repeated learning is thought to be critical for stabilizing learning (Shibata et al., 2017a, 2017b). Follow up studies would be required to determine the optimal training strategy for maximizing long term perceptual gains.

Training listeners to narrow their temporal binding window often decreases their likelihood of integrating auditory and visual stimuli (McGovern et al., 2016; Setti et al., 2014), and the temporal binding window itself is task and stimulus dependent (M. A. De Niear, Gupta, Baum, & Wallace, 2018; Megevand et al., 2013; R. A. Stevenson & Wallace, 2013). Decreased integration can be successfully modelled within a Bayesian causal inference framework as resulting from both an increase in the precision of timing estimates and a decrease in a prior belief that signals originate from the same source (McGovern et al., 2016). Here, we report enhanced auditory visual integration after training listeners to make temporal coherence judgments. Unlike studies that train listeners to narrow their perceptual binding window, in our training paradigm participants were effectively trying to detect small amounts of temporal coherence and distinguish this from fully independent stimuli.

The width of the temporal binding window predicts susceptibility to sound induced flash illusions in naïve listeners (Stevenson, Fister, Barnett, Nidiffer, & Wallace, 2012). We did not find a relationship between the ability of naïve listeners to assess temporal coherence and their ability to exploit temporal coherence between the target and the visual stimulus. Nonetheless, we did observe a correlation between the ability of listeners to discriminate temporal coherence and the relative pattern of performance of the masker-coherent and independent condition with those people who were best able to assess temporal coherence showing a benefit of masker-coherent over independent, and those people who were worse as assessing temporal coherence being relatively impaired on the masker coherent condition relative to the independent condition.

Previous studies have illustrated that visual cues can assist speech processing in noise (Grant, Walden, & Seitz, 1998; Helfer & Freyman, 2005; Schwartz, Berthommier, & Savariaux, 2004). While speech reading abilities are strongly predictive of audiovisual benefit for speech reception thresholds (MacLeod & Summerfield, 1987), lip reading can influence auditory streaming (Devergie, Grimault, Gaudrain, Healy, & Berthommier, 2011), supporting the idea that, in addition to conveying phonetic information, lip reading benefits in noise potentially comprise of both bottom-up sensory effects that facilitate auditory scene analysis (Atilgan et al., 2018) Previous studies exploring the transfer of effects from training on temporal simultaneity judgments to other multisensory paradigms have had mixed results with transfer occurring to some tasks but not others (McGovern et al., 2016; Powers Iii, Hillock-Dunn, & Wallace, 2016; Setti et al., 2014; Sürig et al., 2018). An important question in interpreting the significance of our findings is whether the benefits in the auditory selective attention task transfer to other more real-world tasks such as utilizing speech reading in noisy listening conditions.

### Context

We have previously (Maddox et al., 2015) demonstrated that a temporally coherent visual stimulus can enhance the ability of listeners to focus on one sound in a mixture. Using the same stimuli, we demonstrated a bottom up mechanism through which visual information could change the way in which sound mixtures were represented in auditory cortex, such that the neural representation of sounds that were temporally coherent with a visual stimulus were enhanced. One observation we made from our behavioural data was that listeners varied greatly in their ability to use visual to augment auditory scene analysis. In this study is therefore a first attempt to understand whether we could train listeners to use visual information more effectively. Our longer term goal is to relate these findings to other situations that require focusing on one sound in a mixture, such as listening to speech in noise, to understand whether listeners might benefit from training in order to exploit visual information more effectively.

## Acknowledgments

This work was funded by a Wellcome Trust – Royal Society Sir Henry Dale Fellowship to JKB (ref: 098418/Z/12/Z) and an Action on Hearing Loss PhD studentship to HA. We are grateful to Suganya Mariyanesan for assistance in collecting the control data for this project, and to Ross Maddox and KC Lee for discussion of this work.

